# Inference of annealed protein fitness landscapes with AnnealDCA

**DOI:** 10.1101/2023.05.19.541442

**Authors:** Luca Sesta, Andrea Pagnani, Jorge Fernandez-de-Cossio-Diaz, Guido Uguzzoni

## Abstract

The design of proteins with specific tasks is a major challenge in molecular biology with important diagnostic and therapeutic applications. High-throughput screening methods have been developed to systematically evaluate protein activity, but only a small fraction of possible protein variants can be tested using these techniques. Computational models that explore the sequence space *in-silico* to identify the fittest molecules for a given function are needed to overcome this limitation. In this article, we propose AnnealDCA, a machine-learning framework to learn the protein fitness landscape from sequencing data derived from a broad range of experiments that use selection and sequencing to quantify protein activity. We demonstrate the effectiveness of our method by applying it to antibody Rep-Seq data of immunized mice and screening experiments, assessing the quality of the fitness landscape reconstructions. Our method can be applied to most experimental cases where a population of protein variants undergoes various rounds of selection and sequencing, without relying on the computation of variant enrichment ratios, and thus can be used even in cases of disjoint sequence samples.

## I. INTRODUCTION

The design of proteins to perform a given task (*e.g*. binding a target molecule) is a paramount challenge in molecular biology and has crucial diagnostic and therapeutic applications. Several high-throughput screening technologies have been developed to systematically assess protein activity. Despite the high parallelization of many techniques, a fundamental limitation lies in the small fraction of possible molecules that can be tested compared to the huge number of possible variants. Leveraging those data using effective computational models is crucial to overcome the obstacle by exploring in-silico the sequence space for the fittest molecules for a given function. We use the term *fitness* generically to refer to the protein activity under selection in a screening experiment (or during the *in-vivo* affinity maturation process). Several molecular activities can be selected in such experiments ranging from binding to a substrate to very complex phenotypes, such as conferring antibiotic resistance or multiple unknown interactions in a tissue.

Many machine-learning methods have been proposed recently to learn the protein fitness landscape from sequencing of high-throughput screening experiments [1–7]. Here, we propose a machine learning framework to target sequencing data derived from a broad class of experiments that use selection and sequencing to quantify the activity of protein variants. These experiments include, among others: *Deep Mutational Scanning*, where a library of protein mutants is screened in-vitro for different activities [8–16]; *Directed Evolution* experiments, where a mutagenesis step adds diversity in the library after the rounds of selection [17–19]; sampling of the in-vivo immune response as in antibodies *Repertoire Sequencing* (Rep-Seq) [20]. Some of these experiments serve to select the fittest variants within the screened library while providing quantitative information about the protein activity landscape.

A basic quantitative measure of protein fitness can be obtained by computing the ratio between the relative frequencies of the variants in the populations before and after selection. This ratio, called *selectivity*, is a proxy for the probability that a variant survives the selection process, and has been widely used in the analysis of Deep Mutational Scan experiments [8, 21, 22]. Other approaches leverage more efficiently the same information, by parameterizing in some way the genotype-fitness map [7], or by developing adequate denoising procedures [21, 23].

All these approaches evaluate the fitness from the temporal trajectory of variant abundances through the selection rounds. Conversely, many experimental setups are incompatible with the notion of a single variant trajectory in the population. Such is the case in Directed Evolution experiments, where a mutagenesis process occurs alongside selection that modifies the pool of individual variants from one round to the next [17, 18]. Depending on the interplay between mutational drift, selection strength, and the fitness landscape, the probability to re-sample previously-seen variants can be very small after some rounds. Most variants do not persist through the whole time series and often are observed only once. In other setups, a severely undersampled regime precludes the repeated observation of individual variants. In repertoire sequencing, the coverage is generally too low in comparison to the large number of receptors present in an immune repertoire, which implies that individual sequences are not sampled more than once.

For *in-vitro* screening experiments, factors such as selection strength, number of rounds, the shape of the fitness landscape, the size of the initial library, and sequencing coverage, can limit the ability to observe a relevant fraction of the possible variants. In these cases, we cannot detect the time trajectory of the frequency of most variants and thus we cannot compute an enrichment ratio. Nevertheless, it is still possible to make inferences about the fitness landscape. Another approach involves a dimensionality reduction of the protein sequence space through the modelization of the evolution of the variant distribution as selection proceeds.

Here, we propose AnnealDCA, a simple but effective strategy to perform this task, which can be applied to different experiments and types of data. Our approach is inspired by the simulated annealing method [24] from statistical physics to solve optimization problems. The different experimental rounds can be viewed as an effective cooling process, where an effective temperature is gradually reduced across successive rounds, and the selective pressure becomes increasingly dominant. The general mathematical framework and the associated statistical inference method can be applied to most of the experimental cases where a population of protein variants undergoes different rounds of selection, and a subset (or all) rounds are sequenced. Datasets of this type include, among others, protein screening experiments with one or multiple panning rounds, and the collection of Rep-Seq samples at different infection times.

To demonstrate the effectiveness of our scheme, we apply the method to antibody Rep-Seq data of immunized mice and we predict the antibody affinity towards its cognate antigen. We further test the method in more controlled experiments and assess the quality of the in-silico reconstructed fitness landscape.

## II. METHOD

To describe our method, we start for the sake of simplicity by considering a simple screening experiment of an initial library that takes place over several panning rounds. Other experimental setups will be described next. We define *P*_*t*_(**S**) as the probability of observing a sequence **S** at round *t*. Eventually, *P*_*t*_(**S**) is the quantity we want to estimate from the sequencing data. We introduce a sequence-dependent *survival factor Q*_*t*_(**S**). This quantity is a measure of the probability that sequence *S* survives between round *t −* 1 to *t*. Similarly to [1, 25, 26], we assume that this quantity takes the following exponential form:

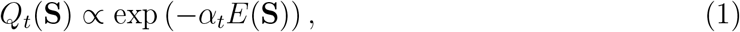

with a time dependent factor *α*_*t*_, modeling the scale of the selective pressure acting at round *t*. The time-independent function *E*(**S**), associates a statistical *energy* to the protein sequence **S**. Thanks to Eq. (1), we can then express *P*_*t*_(**S**) as:

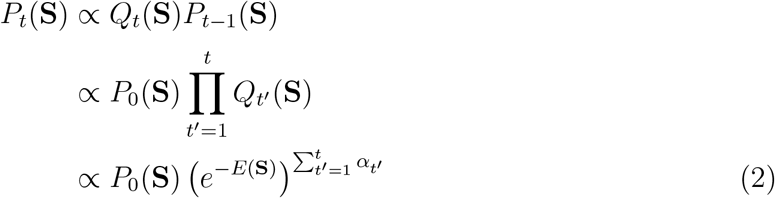

up to a normalization constant. Using Eq. (2), we can express *P*_*t*_(**S**) as a product of the initial configuration probability *P*_0_(**S**) and the factor *e*^*−E*(**S**)^, raised to the sum of the selective pressures of all rounds. We can redefine such a sum as:

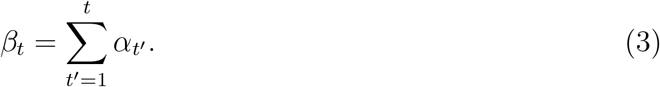

Eq. (3) can be interpreted as a fictitious inverse temperature, accounting for the cumulative selective pressure up to round *t*. In the absence of mutations and if the experimental conditions are the same for all rounds, Fisher’s fundamental theorem of evolution states that *α*_*t*_ is a decreasing function of time [27]. Thanks to Eq. (3), we can trasform Eq. (2) as follows:

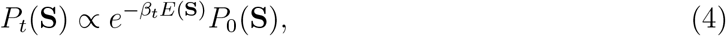

The accumulated selection, quantified by the inverse temperature *β*_*t*_, tends to drive the mass of the distribution *P*_*t*_(**S**) towards the minima of *E*. This mental picture is reminiscent of the simulated annealing process studied in statistical mechanics and other areas [24].

At *t* = 0, *P*_0_(**S**) is the distribution of the variants in the initial library. Since this library is randomly generated, it is supposed to be unrelated to the selection process, and consequently to fitness. We can model the distribution of the initial variants by another similar energy function *G*(**S**):

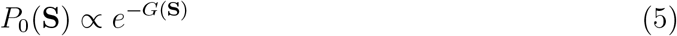

so that Eq. (4) takes the following form:

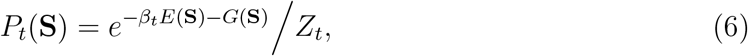

where *Z*_*t*_ = Σ_{S}_ exp (*−β*_*t*_*E*(**S**) *− G*(**S**)) is a time-dependent normalization factor, and the sum runs over all possible sequences.

Fig. 1 shows a pictorial representation of the overall modeling of the experimental screening process. Notably, we do not need any explicit assumption on the specific temporal dependence of the inverse temperature, as the *β* factors can be inferred directly from the data.

**FIG. 1.**
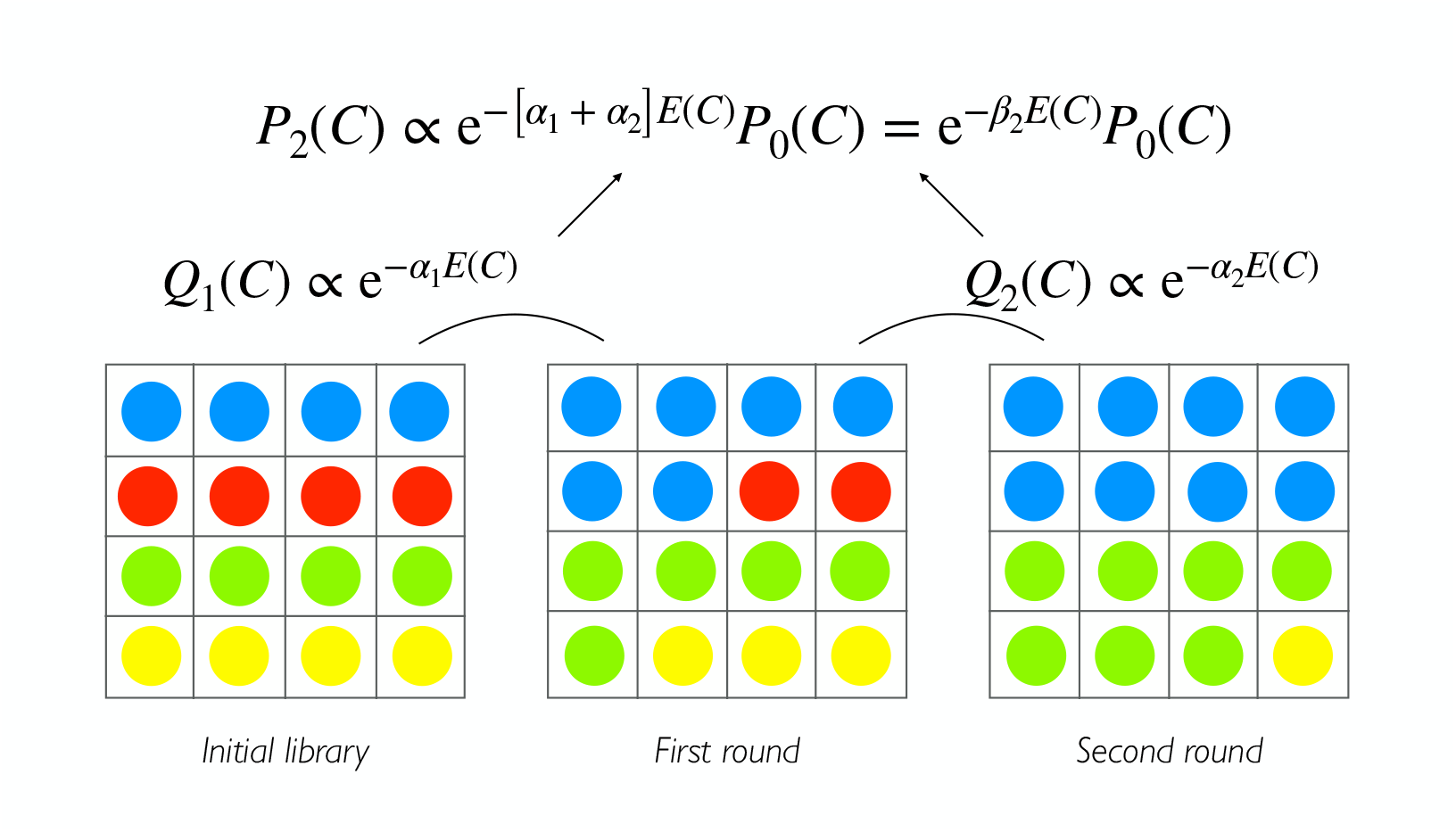
A simplified portrayal of the modeling of the selection process. Each color represents a different variant. Starting from the initial distribution of variants, the probability of observing a sequence in a subsequent round is shaped by the selection process, defined by the energy function *E*(**S**). The selective pressure at each transition is encoded in *α* _*t*_, and the overall fictitious inverse temperature is given by the sum of all transitions 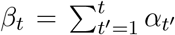 As in a simulated annealing stochastic process where the temperature is progressively lowered, *β*_*t*_ increases with time, thus constraining the variant probability distribution to peak around the fittest sequences.

### Fitness map

The genotype-to-fitness map here is encoded in the energy function *E*. The choice of its functional form and the related number of parameters to be inferred are eventually a trade-off between the expressive power and the actual availability of the sequence data to train the model. One of the simplest parameterizations is an independent site model, where each amino acid contributes additively to the energy:

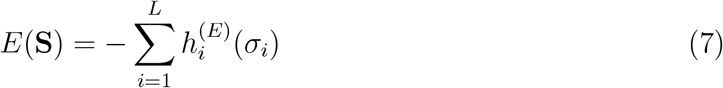

with parameters 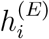 that depend on the identity of the amino acid *σ*_*i*_, present at position *i* along the sequence **S**.

A more complex parameterization is obtained by including pairwise epistatic interactions between all pairs of amino acids and is now widely used in structural biology [28, 29] and functional biology [30–34]. The resulting energy function takes the form of a generalized Potts model:

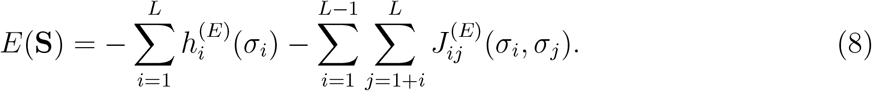

In comparison to the simple independent site model in Eq. (7), the parameterization in Eq. (8) is characterized by 𝒪 (*L*^2^) additional parameters, 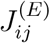, to model the pairwise interactions.

### Model training

The model parameters are trained by maximizing the log-likelihood of the full dataset:

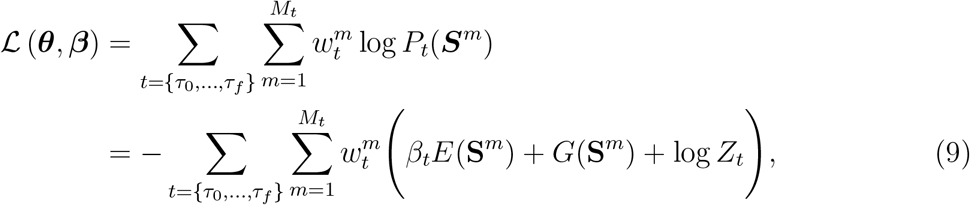

where 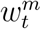 is the normalized abundance of the sequence *m* at time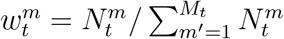, {*τ*_0_, *τ*_1_, …, *τ*_*f*_} ⊂ {0, *t*_1_, …, *t*_*f*_} is the subset of sequenced rounds and ***θ*** = {***θ***^*E*^, ***θ***^*G*^} is the set of parameters of the energy functions.

The likelihood is the product of the probabilities to sample the observed sequences and frequencies from the model at each time point. It can be interpreted as minus the crossentropy between the predicted distribution of the variants and the observed one, the empirical frequencies. The exact maximization of the likelihood involves the computation of the partition function of the model, whose computational complexity scales as 𝒪 (*q*^*L*^). To overcome this practical limitation, there are many approximate methods developed for specific parametrizations of the energy function. Other approaches, based on Monte-Carlo, are more general but might have convergence issues that are difficult to control in practice and are computationally costly. A very effective approach relies on the maximization of a quantity related to the likelihood, called the pseudo-likelihood function [35, 36], whose precise definition, in the case of the Potts model (Eq. (8)), is given in Sec. I A of the SI. A regularization term is added to the pseudo-likelihood to both avoid overfitting and to guarantee a faster time of convergence, since the pseudo-likelihood tends to have a large number of flat directions in parameters space. While Eq. (9) is separately convex with respect to the energetic parameters and the inverse temperatures, this is no longer true when both are inferred simultaneously. To learn the parameters 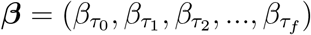 it is possible to use an iterative optimization scheme: starting from an arbitrary set of ***β*** components, the energetic parameters ***θ*** are optimized. Next, the ***β*** components are updated while the ***θ*** are kept fixed. The two steps are iterated until convergence of both sets of parameters is reached. Some constraints can be imposed on the *β* parameters without affecting the expressivity of the model. In particular, it is possible to set 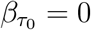 and to fix a scale factor setting 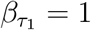.

## III. RESULTS

Most of the computational methods used to infer the fitness landscape from screening experiments rely on the computation of the enrichment/depletion ratios for a sufficiently large set of variants to train a regression model. The enrichment ratio is, in its simpler form, the ratio between the frequency of a variant in different rounds (see Eq. (4) in the SI). This quantity is a proxy for the ability of a variant to be selected during the process, namely, the fitness. However, several cases exist in which the temporal trajectory of the single variant is not detected. It can happen when: (i) the experiment is dominated by noise effects; (ii) the sequencing coverage is not adequate in comparison to the broadness of the library, and under-sampling effects might dominate; (iii) some mutations are introduced along the selection process at each round of the experiments. As a consequence most of the variants sampled at different time points could be unique or in low copies, affecting the accuracy of the enrichment ratios estimate. Conversely, the generality of our approach makes it applicable to all the above cases. We demonstrate the efficacy and the versatility of the method by applying it to three different experimental setups (the experimental datasets are listed in Tab. I of the SI), which are described briefly below:

- *Antibody Repertoire Sequencing (Rep-Seq)* The Antibody Repertoire refers to the set of functionally diverse immunoglobulins in an individual at a given time. The Rep-Seq technique allows for studying a sample of the immunoglobulin repertoire distribution. We collect the data from Khan et al. [37] and Gerard et al. [38]. In the first paper, the authors isolated and sequenced the IgG secreted by memory B cells and plasmablasts of not-immunized mice, whereas in the second the same mouse clones were immunized against an antigen. Then, the isolated IgG repertoire undergoes a high-throughput phenotypic assay with a microfluidic platform whose output is enriched in antigen binders.
- *Deep mutational scanning (DMS)* These experiments combine high-throughput screening of a mutational library with sequencing to assess the effect of mutations on protein activity [39]. An initial library of protein variants undergoes one or multiple cycles of selection for a protein function (*e.g*. binding to a substrate). After a number of rounds, a sample of the variants population is deep-sequenced to assess the variants’ abundances and their changes over time. Typical examples are *in-vitro* display experiments (*e.g*. phage display). The experiments and datasets we used in our study are described in Fowler et al. [22], Boyer et al. [40], and Wu et al. [41].
- *Directed Evolution experiments (DE)* DE experiments follow a setup similar to DMS, with the difference that in this case, random mutations are repeatedly introduced before each panning round. In some cases, the experiment starts from a single wild-type protein. The experiment attempts to simulate *in-vitro* a natural Darwinian evolutionary process, where mutations explore the sequence space creating new genotypes whose phenotype is tested for the protein function. The experiments and datasets are described in the following two papers: Fantini et al. [17], Stiffler et al. [18].

### A. Antibodies Repertoire Sequencing

By applying the method to antibodies Rep-Seq data, we want to infer the probability of an antibody being the outcome of an immune response. Once the model is trained, we obtain a parameterization of the probability function that can be used to design novel antibodies.

As the immune system reacts to some antigen, in principle, the designed antibodies should have a high affinity to the same antigen. We use two datasets for this purpose: a sample of mice unimmunized repertoires, which we name background or negative dataset and a sample of the immunized repertoire (positive set). In this case, the sample after the immune response is enriched in binders by functional sorting through a microfluidic platform. In general, we could also directly employ some samples from a repertoire (see Asti et al. [30]).

The general idea is to model the probability of observing an antibody in the positive set as the product of the following two probabilities [25, 26]: The background probability, *i.e*. the probability to observe an antibody in the negative set (the unimmunized repertoire), and the selection factor that describes the overall effective process of the immune response (together with the enrichment platform in this case), see Fig. 2 for a pictorial representation. The negative or background dataset contains sequences from the IgG heavy chain repertoire of three unimmunized BALB/c mice (the same type as for the positive dataset). The dataset is publicly available from the Observed Antibody Space [42] and the experimental setup is described in Khan et al. [37]. The dataset contains a total of 19772 unique IgG heavy chain sequences with the number of readouts. The positive datasets contain sequences of IgG heavy chain (VH) of immunized BALB/c mice repertoire sorted by a droplet microfluidics platform by the binding status of two immunogenic targets: Tetanus toxoid (TT) and Glucose-6-Phosphate Isomerase (GPI). The number of unique VH sequences is 3881 for TT and 3233 for GPI. All sequences were aligned using the Martin antibody numbering. The complete preprocessing pipeline is described in the supplementary material.

**FIG. 2.**
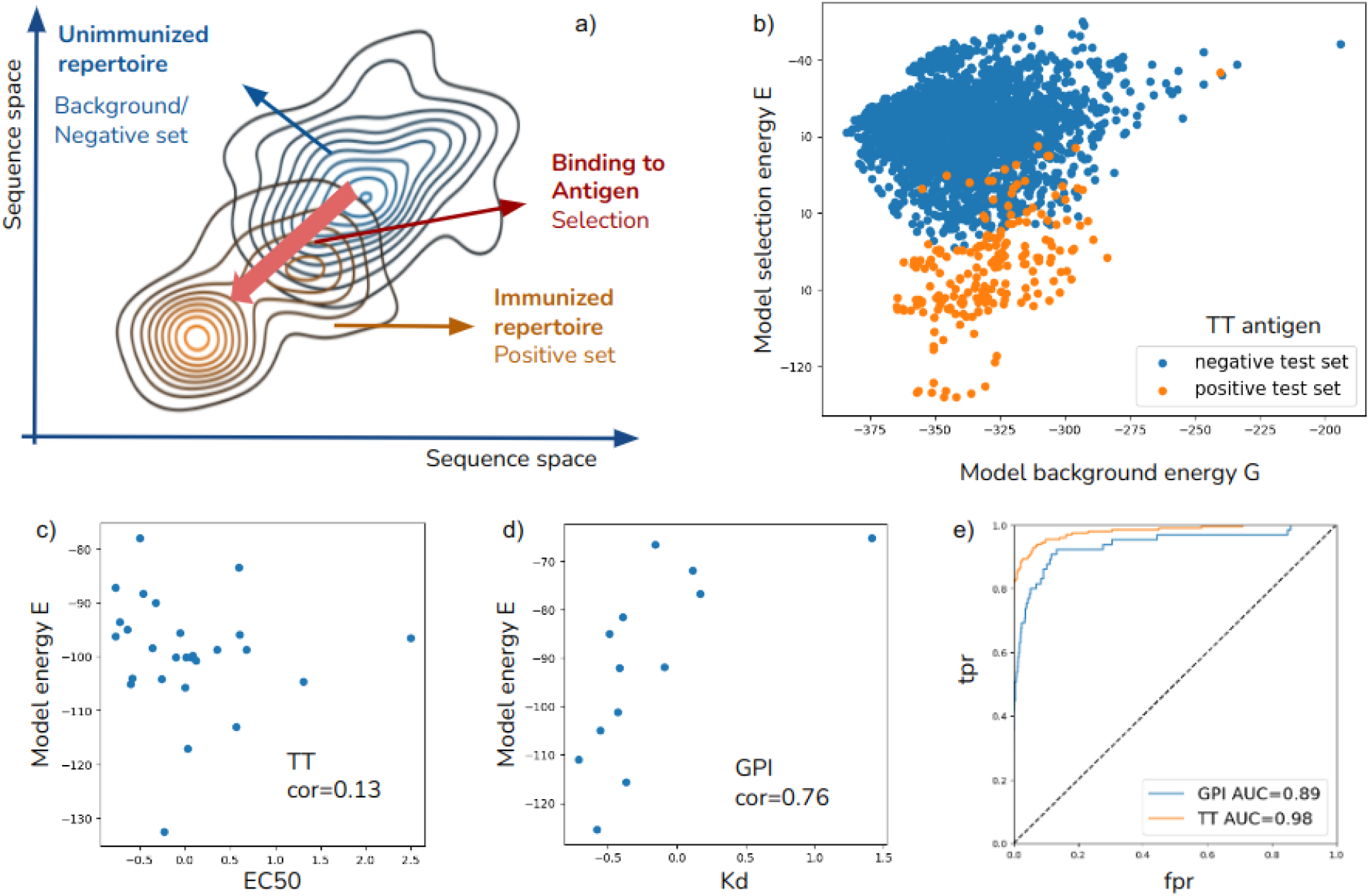
Application of the method to antibody Repertoire Sequencing data. In panel (a), a representation of the inference of the selection process: the initial library is the unimmunized repertoire (negative set), while after the immune response, the library is shaped by the selection to bind the antigen (positive set). In panel (b), the plot of the background and the selection energy for negative and positive sets shows that the selection energy separates the antibodies of the immunized repertoire from the unimmunized one. The capacity of the model to discriminate binders and not-binders can be assessed as the classification on a test set composed of negative and positive antibodies. Panel (c) shows the ROC curves for the GPI and TT cases, demonstrating almost perfect classification with the area under the curve (AUC) of 0.89 and 0.98 respectively. In panels (d) and (e), the scatter plots display the comparison of the selection energy and the affinity measures for a set of antibodies: EC50 values for TT and *K*_*d*_ measures for GPI. Only in the second case, there is a significant correlation (Spearman coefficient 0.76) demonstrating the capacity of AnnealDCA to learn the binding affinity towards the antigen from repertoire sequencing data.

We analyzed the similarity of positive and negative datasets in terms of different sequence statistics such as distances from consensus sequence, site conservation and covariance, alignment PCA, and germline distributions. These preliminary analyses however were not capable of revealing sensible differences between the two datasets (see more details in Sec. II A and Fig. 1 of the SI).

We then proceed to test the ability of the model energy to discriminate between binders and not binders. For this purpose, we split the positive and background datasets into a training set and a test set to validate the classification predictions. This validation procedure was chosen due to the lack of a large list of IgG labeled as binders of the two antigens (TT and GPI). Thus, we use the positive and background set as a proxy for binders/not binders labels. Fig. 2 shows the results of the classification task. The sequences with low background energy are likely to be present in the unimmunized repertoire, while the selection energy accounts for the probability of an antibody being part of the immune response to the specific antigen. The model can discriminate remarkably well the binders (of the positive set) and not binders (of the background set), as demonstrated by the ROC curve in the test sets of both targets (AUC 0.98 for TT and 0.89 for GPI).

In Gerard et al. [38], the authors reported the experimental measures of the dissociation constant *K*_*d*_ with GPI of a small set of antibodies (14 binders and 2 not-binders) and the EC50 values against TT for another small set (42 binders and 4 not-binders). Using this validation set, we can test whether the inferred selection energy correlates with the antibody affinity or the neutralization power. The antibodies in the validation test are sampled fairly in the sequence space from the positive ensemble and lay on the high-selectivity model energy region. The results show that inferred selection energy correlates with *K*_*d*_ GPI measures, while there is no significant correlation between selection energy and EC50 in the TT case (see Fig. 2). The reason could be that the microfluidic platform sorts the antibodies based on binding to antigen while EC50 measures antigen neutralization (the half maximal effective concentration of Abs).

### B. Deep mutational scanning (DMS)

Deep mutational scanning experiments are explicitly designed to quantify mutation effects on fitness. The broadness of the library and the sequencing depth are chosen to compute reliable enrichment measures for the variants ([8, 22]). Thus, approaches that leverage the enrichment ratios are more suitable to address these datasets. Nevertheless, it provides an interesting controlled case to assess the inference procedure and compare it to other tools. The screening experiment described in [22] probes the binding affinity of the human WW-domain with its peptide ligand. More than 6 × 10^5^ unique variants are generated in the initial library, which comprises almost all single point mutations, a fourth of the double mutations, and almost 2% of all three point mutations. Then, six rounds of phage display screening are performed, and rounds 3 and 6 are sequenced. In Boyer et al. [40], the library variability leans on a short sequence segment of the CDR3 region of an antibody’s heavy chain (*L*=4). The library (chosen among 24 different scaffolds around the CDR3 region) is subsequently screened for three rounds of panning against a polyvinylpyrrolidone target.

In the experiment described in Wu et al. [41], the variants library contains all possible mutations of four residues of the IgG-binding domain (GB1). The library is then screened to bind an immunoglobulin fragment target in a single round of selection.

We perform a 5-fold cross-validation to test the inference method: for each dataset, a random selection of 4*/*5 serves to train the model while 1*/*5 operates as a benchmark. As in [7], we compare the model energy function with the log-selectivity *θ* ^*m*^ (see Eq. (4) in the SI), which is computed from the enrichment ratios and serves as a proxy for the variants fitness. The performance is then evaluated by the Pearson correlation between the model energies *E* and the log-selectivities in the test set. On all datasets, we observe high correlation as shown in panel (a) of Fig. 3. Finally, we compare the results with the inference method developed in [7]. As expected, the [7] model performs better in all three datasets, as it uses the enrichment information. Nevertheless, we underline that we are still able to obtain an energy function highly correlated with fitness, close to the best performance. Furthermore, for the Wu et al. dataset [41], we notice how the discrepancy between the two methods becomes very shallow, due to the broad coverage of sequence space. Lastly, we remark that [7] is unable to handle other datasets considered in this work.

**FIG. 3.**
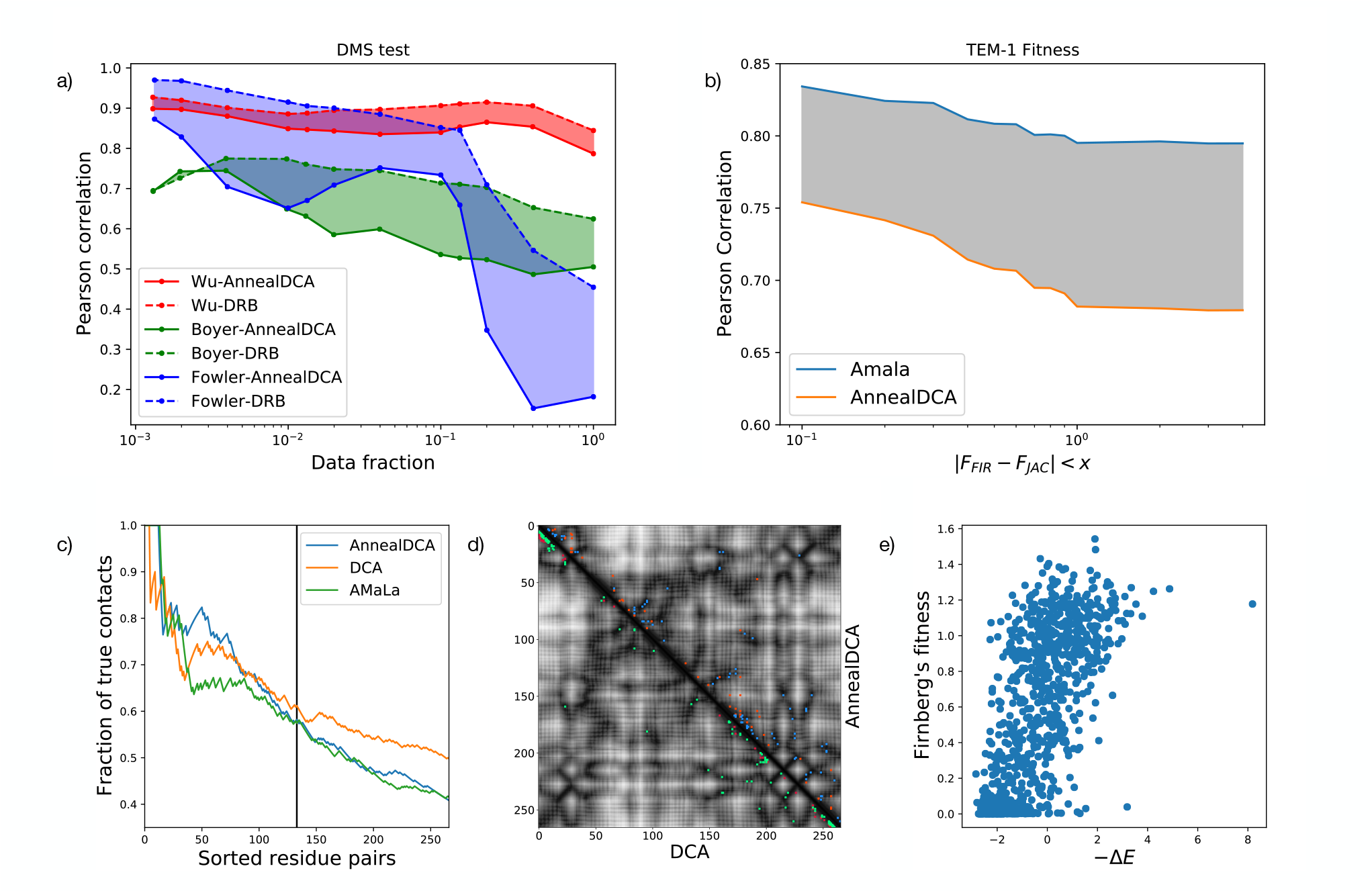
Results of the method application to Deep Mutational Scanning [22, 40, 41] and Directed Evolution [17, 18] experiments. Panel (a) shows the overall performance of the method on DMS: the Pearson correlation coefficient between inferred selective energies *E* and empirical log-selectivities (Eq. (5) of the SI). The correlations are reported as a function of data fraction pruned for the level of noise (error in the regression of empirical selectivities, as in [7]). Results are compared with the method described in [7], which gains by using the enrichment information. Panel (b): comparison between AnnealDCA and AMaLa [6] for the reconstruction of the fitness landscape of TEM-1 from [17] data. Accuracy is quantified via the Pearson correlation between inferred energies *E* and independent fitness measurements [15, 16], as a function of the threshold discrepancy between the two datasets *x*. Panel (c): contact prediction sensitivity plot for the protein PSE-1. AnnealDCA, AMaLa, and plmDCA are inferred using data of [18] (DCA uses the last round only, as in [5]). On the y-axis, the *positive predicted value* (PPV) is reported as a function of the residue pairs, sorted in decreasing order of the Frobenius norm (see Sec. I C in the SI). Panel (d): Contact map related to panel (c). The plot is an *L*×*L* representation of the possible contacts between protein residues. The prediction of DCA (lower-left) and AnnealDCA (upper-right) are compared; correctly/incorrectly predicted contacts are respectively reported in green/red for DCA and blue/orange for AnnealDCA. Panel (e): scatter plot between selective energies inferred on [17] and fitness measurements of [16] for a specific threshold value *x* = 0.8. Energies are rescaled with respect to the wild-type sequence Δ*E* = *E*(**S**) *− E*(**S**^*wt*^).

### C. Directed Evolution experiments

Due to its flexibility, we can apply the method also to experiments in which new protein variants appear via a mutagenesis step at each new round. In this case, as discussed in [6], we cannot compute the enrichment ratios and selectivity. The *G* Hamiltonian in Eq. (6), although being time-independent, accounts for the mutational step in an effective manner, as is demonstrated by the high correlation between *G* and the Hamming distance from the wild-type sequence (see Fig. 6 of the SI). The *E* part, on the other hand, corresponds to the selection process.

We collect data from three experiments described in Fantini et al. and Stiffler et al. [17, 18]. The authors screen proteins responsible for antibiotic resistance in bacteria: TEM-1 and PSE-1 variants of the *β*-lactamase family and AAC6 protein of the acetyltransferase family. Starting from a wild-type protein, error-prone PCR creates new mutants at each round. Subsequently, the library undergoes a selection step in which bacteria equipped with the mutants are exposed to an antibiotic-rich environment. This cycle of mutagenesis and screening is repeated multiple times and for a subset of the panning rounds a sample of the library is sequenced. Specifically, 20 rounds of Directed Evolution at an ampicillin concentration of 6*μ*g/mL are performed for PSE-1, among which rounds 10 and 20 are sequenced, whereas AAC6 mutants are subjected to 8 rounds at a kanamycin concentration of 10*μ*g/mL, of which rounds 2, 4 and 8 are sequenced. Finally, in [17] TEM-1 mutants are exposed to two different antibiotics concentrations: 25*μ*g/mL for all rounds but 5 and 12, for which the concentration is raised to 100*μ*g/mL. Out of the 12 experimental cycles, rounds 1, 5, and 12 are sequenced.

We performed two different validations to assess the inferred model:

i. In the case of TEM-1 *β*-lactamase, we directly compare the model energy with independent fitness measurements related to antibiotic resistance, collected from [15, 16]. In [15] variants fitness is quantified in terms of *minimum inhibitory concentration* (MIC), that is, the minimum antibiotic concentration necessary to neutralize bacteria equipped with that variant. On the other hand, in [16], the authors directly measured the gene fitness (see Sec. I D of the SI). For our analysis, we mapped the measurements of [16] onto those of [15], following the procedure outlined in [31]. The results show that the inferred energy correlates with the experimental fitness (see panel (b) of Fig. 3). The method described in [6], specifically designed for these experiments performs systematically better.
ii. In the PSE-1 and AAC6 cases, for which fitness measurements are not available, we validate the model using the prediction of protein structure contact map as prescribed by the DCA method [28, 29]. Then, the predictions are compared to the crystallographic studies of the protein structures. The contact predictions are obtained using the coupling parameters **J**, which quantify the interaction between residues in the DCA framework [28, 29]. We used the Frobenius norm of the *q* × *q* matrix *J*_*ij*_ to obtain a score quantifying the epistatic interactions between pairs of positions (see Eq. (5) in SI), on top of which we apply the *average product correction*. These residues are more likely to be found in spatial proximity in the folded structure as shown in panel (c)(e) of Fig. 3.

## IV. CONCLUSIONS

Several machine-learning methods have been proposed for learning a protein fitness landscape using sequencing data obtained from high-throughput screening experiments [2, 7, 25, 26]. However, these methods require observation of the trajectory in multiple rounds of selection of a statistically relevant set of variants. This presents a limitation as detecting the single variants trajectory in the population is often unfeasible in many experimental setups. To overcome this issue, we propose AnnealDCA, a novel machine-learning framework inspired by the simulated annealing method from statistical physics [24]. This approach can handle sequencing data derived from a broad range of experiments that use selection and sequencing to quantify the activity of protein variants, including Deep Mutational Scanning, Directed Evolution experiments, and antibodies Repertoire Sequencing (Rep-Seq), among others.

In our approach, the selection process acts as a cooling process where the distribution of the population on the fitness landscape is gradually peaked around regions of higher fitness. The samples before and after the selection are considered at different statistical temperatures and the inference method decouples the distribution contribution due to the initial library and the time-dependent fitness part. The general mathematical framework and the inference method can be applied to most of the experimental cases where a population of protein variants undergoes a selective process and is sequenced at different times. Such datasets include, among others, protein screening experiments with one or multiple panning rounds, and the collection of Rep-Seq samples at different infection times.

To demonstrate the effectiveness of this approach, we applied the method to antibodies Rep-Seq data of immunized mice to predict the antibody’s affinity towards the antigen. We learned the model energies from the repertoire of mice unimmunized and subjected to two antigens. The model energy was then used to accurately classify binders and not-binders to the antigen. This was supported by the fact that it correlated well with experimental measures of the *K*_*d*_ antibody-antigen of a set of antibodies not used in the training of the model.

To further test our approach, we applied it to more controlled experimental setups using three Deep Mutational Scans experiments. The results of 5-fold cross-validation showed a high correlation between the inferred fitness landscape and the experimental selectivity. Additionally, we applied the method to Directed Evolution experiments of three proteins responsible for antibiotic resistance in bacteria, where mutations are added to increase the variability and explore sequence space around a wild-type sequence. The model energy precisely described the antibiotic resistance measurements of a set of variants. Moreover, the model couplings parameters were able to predict the three-dimensional contact map with a level of precision comparable to other approaches.

In summary, AnnealDCA provides a simple but effective strategy that can be applied to different experiments and data types where a population of protein variants undergoes a selective process and is sequenced at different times.

## V. DATA AND CODE AVAILABILITY

The validation data used in this study, the models and the script for training the model (coded in Julia language), the Jupyter notebooks to retrain the models and redo the figures, and the data preprocessing scripts are available on Gitlab: https://gitlab.com/luca.sesta/AnnealDCA.jl.

## ACKNOWLEDGMENTS

AP acknowledges funding by the EU H2020 research and innovation programme MSCA-RISE-2016 under Grant Agreement No. 734439 INFERNET, as well as for financial support from FAIR (Future Artificial Intelligence Research) and ICSC (Centro Nazionale di Ricerca in High-Performance Computing, Big Data, and Quantum Computing) founded by European Union – NextGenerationEU. We thank Adam Woolfe, Andreas Raue and Annabelle Gerard for helpful discussions and assistance with the Rep-seq data.

