## Supplementary materials for "Inference of annealed protein fitness landscapes with AnnealDCA"

[5] *Co-last author*

**CONTENTS**

I. Methods 2

A. Pseudo-likelihood 2

B. Selectivity 3

C. Contact prediction 4

D. TEM-1 fitness measurements 4

II. Datasets 5

A. Antibodies Repertoire Sequencing 5

B. Deep Mutational Scanning 6

C. Directed Evolution 7

III. Further results 8

A. Deep Mutational Scanning 8

B. Directed Evolution 8

References 8

### I. METHODS

#### A. Pseudo-likelihood

Direct optimization of the objective function in Eq. (9) of the main text is infeasible due to the necessity to compute the partition function. To work around this limitation, we employed the so called *pseudo-likelihood* approximation [1, 2], which is based on single-site conditional probabilities. For the AnnealDCA model, such probabilities read:

$$P_t(\sigma_r|\boldsymbol{\sigma}_{\setminus r}) = \frac{\exp\left\{\beta_t\left[h_r^{(E)}(\sigma_r) + \sum_{i\neq r} J_{ri}^{(E)}(\sigma_r,\sigma_i) + h_r^{(G)}(\sigma_r) + \sum_{i\neq r} J_{ri}^{(G)}(\sigma_r,\sigma_i)\right]\right\}}{\sum_{a=1}^{q}\exp\left\{\beta_t\left[h_r^{(E)}(a) + \sum_{i\neq r} J_{ri}^{(E)}(a,\sigma_i) + h_r^{(G)}(a) + \sum_{i\neq r} J_{ri}^{(G)}(a,\sigma_i)\right]\right\}}. \quad (1)$$

The advantage brought about by Eq. (1) is that the normalization factor can be computed as a sum over only $q$ configurations. The objective function we ultimately optimize in order to infer AnnealDCA parameters becomes in this framework:

$$
\begin{aligned}
g\left(\boldsymbol{\theta}^{E}, \boldsymbol{\theta}^{G}, \boldsymbol{\beta}\right) = &-\sum_{r=1}^{L} \sum_{t=\{\tau_1,\dots,\tau_f\}} \sum_{m=1}^{M_t} w_t^m \log P(\sigma_r = \sigma_r^m | \boldsymbol{\sigma}_{\backslash r} = \boldsymbol{\sigma}_{\backslash r}^m) \\
&+ \sum_{r=1}^{L} R_r^{(E)}\left(\mathbf{h}_r^{(E)}, \mathbf{J}_r^{(E)}\right) + \sum_{r=1}^{L} R_r^{(G)}\left(\mathbf{h}_r^{(G)}, \mathbf{J}_r^{(G)}\right) \\
&= \sum_{r=1}^{L} g_r\left(\boldsymbol{\theta}_r^{E}, \boldsymbol{\theta}_r^{G}, \boldsymbol{\beta}\right), \qquad (2)
\end{aligned}
$$

where we introduced the regularization contribution, which we choose to be an $l_2$ penalty:

$$
R_r\left(\mathbf{h}_r, \mathbf{J}_r\right) = \lambda_h \|\mathbf{h}_r\|_2^2 + \lambda_J \|\mathbf{J}_r\|_2^2 = \lambda_h \sum_{a=1}^{q} h_r^2(a) + \lambda_J \sum_{i \neq r} \sum_{a,b=1}^{q} J_{ri}^2(a,b), \qquad (3)
$$

depending on the auxiliary parameters $\lambda_h$ and $\lambda_J$, which set the strength of the regularization. The $l_2$ regularization can be also intepreted as a Gaussian prior over the parameters in a Bayesian framework. Another significant advantage of the pseudo-likelihood approach is that the single site contributions $g_r$ in Eq. (2) can be optimized independently [2], and thus the parameters learning can be easily parallelized. The drawback of such asymmetric approach, is that it provides two different estimates for each coupling parameters $J_{ij}$: $J_{ij}^{i}$ and $J_{ji}^{j}$, depending on whether the parameter was estimated from single site objective $g_i$ or $g_j$. The issue can be solved by combining together the two estimates, defining a unique coupling $J_{ij} = \left(J_{ij}^{i} + J_{ji}^{j}\right)/2$.

##### B. Selectivity

Enrichment ratios, which are defined as the number of copies of a specific sequence $N_t^m$ at a certain time $t$ divided by the number of copies at the previous time $N^{(m,t-1)}$ are often used to quantify fitness in protein screening experiments. From this, it possible to define the log-selectivity according to:

$$
\log\left[\frac{N_t^m}{N_{t-1}^m}\right] = \theta^m + \alpha_t^m + \epsilon_t^m, \qquad (4)
$$

where $\alpha_t^m$ and $\epsilon_t^m$ are respectively an amplification factor and a noise contribution, and $\theta^m$ represents the log-selectivity of sequence $m$.

##### C. Contact prediction

In order to assess contacts prediction between protein residues, we use the Frobenius norm of the $q \times q$ matrix $J_{ij}$ to obtain a score depending on pairs of positions:

$$F_{ij} = \|J_{ij}\|_2 = \sqrt{\sum_{a,b=1}^{q} J_{ij}(a,b)^2}. \tag{5}$$

The Frobenius score is subsequently modified according to the average product correction $F_{ij}^{\mathrm{APC}} = F_{ij} - F_{i\cdot}F_{\cdot j}/F_{\cdot\cdot}$, where the dots represent averages over the corresponding indeces. Residue pairs characterized by a higher value of $F_{ij}$ have higher epistatic interaction and are more likely to be found in spatial proximity in the folded structure, as showed in panel (c)(d) of Fig. 3 in the main text. Specifically, a pair of residues is considered to be in contact if at least two heavy atoms within the residues have a distance less than $8\mathring{A}$. Moreover, we restrict ourselves to consider pairs of residues $i$ and $j$ such that $|i-j| > 4$.

##### D. TEM-1 fitness measurements

In Sec. III of the main text we tested the capability of AnnealDCA to infer the fitness landscape of TEM-1 $\beta$-lactamase from directed evolution experiments data [3]. In order to assess the meaningfulness of the inferred landscape we relied on the experimental fitness measurements realized in [4]. Since TEM-1 is the protein providing bacteria with antibiotics resistance, the mutants fitness will be quantified by a phenotypic trait related to the response to the antibiotics. Specifically, in [4] the authors define the fitness of a TEM-1 mutant as a weighted average of antibiotic concentrations:

$$f_m = \frac{\sum_{p=1}^{13} c_p^m \log_2(a_p)}{\sum_{p=1}^{13} c_p^m}, \tag{6}$$

where $p$ is the index running over a collection of plates characterized by a different antibiotics concentration $a_p$, coinciding in the present case with ampicillin. The weights of the average are provided by $c_p^m$, that corresponds to the number of copies of mutant $m$ in the $p$-th plate. Specifically, Eq. (6) provides the unnormalized protein fitness, whose normalized version is given by $\phi_m = 2^{f_m}/2^{f_{wt}}$, where $f_{wt}$ is the fitness of the wild-type sequence.

#### II. DATASETS

On GitHub at `https://gitlab.com/luca.sesta/AnnealDCA.jl` are available all the data used in this study and the codes to obtain them, that includes the repository of the AnnealDCA method to train the models, the Jupyter notebooks to reproduce the results and the script for the data preprocessing.

| Article | Experiment | Protein | # samples |
|---|---|---|---|
| Khan et al. (2016) | Rep-Seq | IgGHV | 1 |
| Gerard et al. (2020) | Rep-Seq & sort | IgGHV | 2 (TT and GPI) |
| Boyer et al. (2016) | DMS | Ab IgH | 3 (round 1-3-6) |
| Wu et al. (2016) | DMS | GB1 | 2 (round 1-2) |
| Fowler et al. (2010) | DMS | WW | 3 (round1-3-6) |
| Fantini et al (2019) | DE | TEM-1 | 3 (round 1-5-12 |
| Stiffler et al. (2020) | DE | PSE-1 | 2 (round 10-20) |
| Stiffler et al (2020) | DE | AAC6 | 3 (round 2-4-8) |

TABLE I. Table list of datasets used and the related literature.

##### A. Antibodies Repertoire Sequencing

The data are described in Gerard et al. [5] and T.A.Khan et al. [6], the datasets publicly available and can be downloaded from `https://static-content.springer.com/esm/art%3A10.1038%2Fs41587-020-0466-7/MediaObjects/41587_2020_466_MOESM5_ESM.xlsx` and `http://www.ncbi.nlm.nih.gov/bioproject/311999` in the Raw FASTQ data format.

Once translated in fasta format ( conversion script available on github repository) the sequences were align to martin numbering using ANARCI tool `https://github.com/oxpig/ANARCI` [7]. Then, we filter out the aligned positions with more than 98% gap or 99% conserved residues. The full preprocessing piepline are available on Github.

In Fig. 1, the difference of the positive and negative datasets in terms of hamming distances from consensus sequence and the projections over the two first principal components are depicted, showing the overall high similarity of the positive and negative datasets.

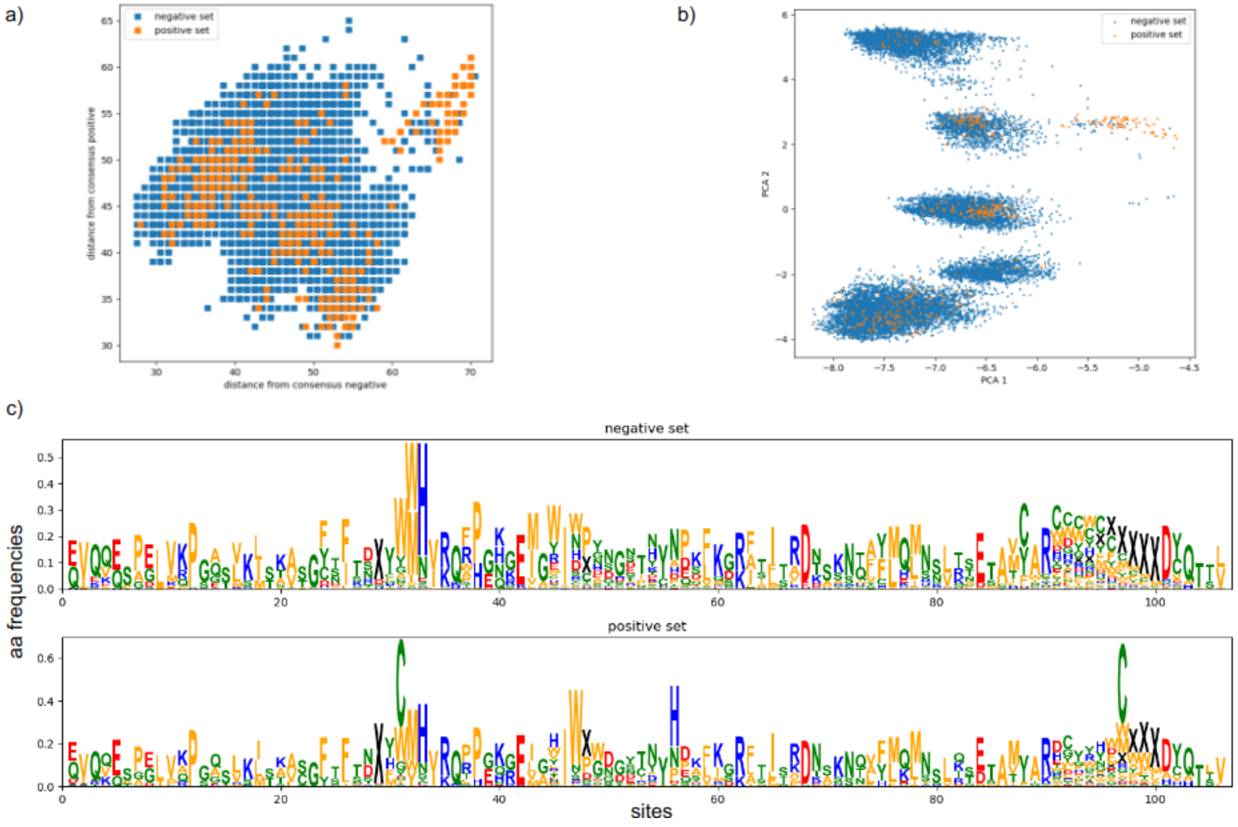


FIG. 1. Antibodies repertoire sequencing data. In this figure three different ways to compare the background or negative sequence set (Khan et al.) and the positive set (Gerard et al. GPI antigen) are shown. In panel (a) the distance of each sequence in both datasets (negative are blue dots and positive orange dots) from the negative consensus sequence (x-axis) and positive consensus sequence (y-axis). In panel b the multi-sequencing alignment (MSA) of the negative dataset is used to compute eigenvectors. The sequences in the two datasets are then projected onto the first two components. In panel (c) the sequence logos of the two datasets is shown.

#### B. Deep Mutational Scanning

The genome sequence data of Wu et al. [8] were downloaded from `https://www.ebi.ac.uk/ena/data/view/SRX958008`. For each DNA sequence a quality score and a pair of forward and backward reads are available. Low quality reads are discarded from the dataset, and unique sequences are constructed from the forward and backward reads in such a way that, if a mismatch between two sites is present, the nucleotide with the higher quality score is retained. A similar procedure is followed for the data of Fowler et al. [9], whose raw genome sequences are downloaded from https://www.ncbi.nlm.nih.gov/sra/SRA020603. In [10], the authors study the binding capability of antibodies onto two targets, namely a short 9 nucleotides DNA segment and synthetic polymer named polyvinylpyrrolidone (PVP). Sequence variability is introduced in the short CDR3 segment, whereas different $V_H$ scaffolds are used to define 24 different libraries. For our analysis, we focused on library F3, in which binding onto PVP is probed. Such data were downloaded from https://www.pnas.org/doi/suppl/10.1073/pnas.1517813113/suppl_file/.

For all DMS datasets, genome sequences are subsequently converted to protein sequences, from which the input data of the method are constructed. Specifically, the aligned unique sequences are concatenated to build up an $L \times M$ alignment matrix Z, with $M$ the number of unique sequences and $L$ the length of the proteins. Associated to this, an $M \times T$ matrix of counts is constructed, with $T$ the number of available sequenced rounds of the experiment (usually including the initial library). In doing so, the sequences characterized by a high uncertainty value on the log-selectivity (Eq. (4)) are filtered out.

##### C. Directed Evolution

The Directed Evolution data considered in this work are described in [3, 11]. Raw data related to [3] are available at the National Centre for Biotechnology Information Sequence Read Archive (SRA), therein accessible via the code PRJNA528665 (http://www.ncbi.nlm.nih.gov/sra/PRJNA528665). More refined data can be found at BioSNS site: http://laborator-iobiologia.sns.it/supplementary-mbe-2019/. Raw data related to [11] can also be found in the SRA, with accession code PRJNA578762, whereas refined data can be downloaded from https://github.com/sanderlab/3Dseq.

Similarly to what we did for the DMS data, the DNA sequences are firstly translated to amino acid ones. From these, an $L \times M$ alignment matrix $Z$ and an $M \times T$ counts matrix are defined. Here, $M$ coincides with the total number of unique sequences produced during the whole experiment, and $T$ corresponds to the number of sequenced rounds.

#### III. FURTHER RESULTS

##### A. Deep Mutational Scanning

In Fig. 2 the scatter plots between the minus selective energy $E$ and the empirical log-selectivities as defined from Eq. (4) are shown for all three experiments [8, 10, 12].

In Fig. 3 we report the analogous of panel (a) of Fig. 3 in the main text, realized over the train datasets.

##### B. Directed Evolution

In panels (c) and (d) of Fig. 3 of the main text we reported respectively the sensitivity plot and the contact map related to the experiment on PSE1 of [11]. Here we show the analogous results obtained for the AAC6 protein. In Fig. 4 we report the sensitivity plots for AnnealDCA, AMaLa and PlmDCA. The first provides a positively predicted value at half-length of $\text{PPV}(L/2) = 0.51$ and an *area under the curve* (AUC) of $\text{AUC}(L/2) = 0.54$, comparable with the results provided by the AMaLa method ($\text{PPV}(L/2) = 0.51$, $\text{AUC}(L/2) = 0.51$) and significantly better than PlmDCA ($\text{PPV}(L/2) = 0.31$, $\text{AUC}(L/2) = 0.34$). In Fig. 5 we show the contact maps obtained with the AnnealDCA and the DCA methods.

Finally, in Fig. 6 we show a scatter plot between AnnealDCA energies $G$ computed over a set of sequences from the AAC6 experiment [11] and their corresponding Hamming distance from the wild-type sequence. The two quantities turn out to be highly correlated, as their Pearson correlation coefficient is 0.94.


[1] M. Ekeberg, C. Lövkvist, Y. Lan, M. Weigt, and E. Aurell, Improved contact prediction in proteins: using pseudolikelihoods to infer potts models, Physical Review E **87**, 012707 (2013).

[2] M. Ekeberg, T. Hartonen, and E. Aurell, Fast pseudolikelihood maximization for direct-coupling analysis of protein structure from many homologous amino-acid sequences, Journal of Computational Physics **276**, 341 (2014).

[3] M. Fantini, S. Lisi, P. De Los Rios, A. Cattaneo, and A. Pastore, Protein Structural Informa-

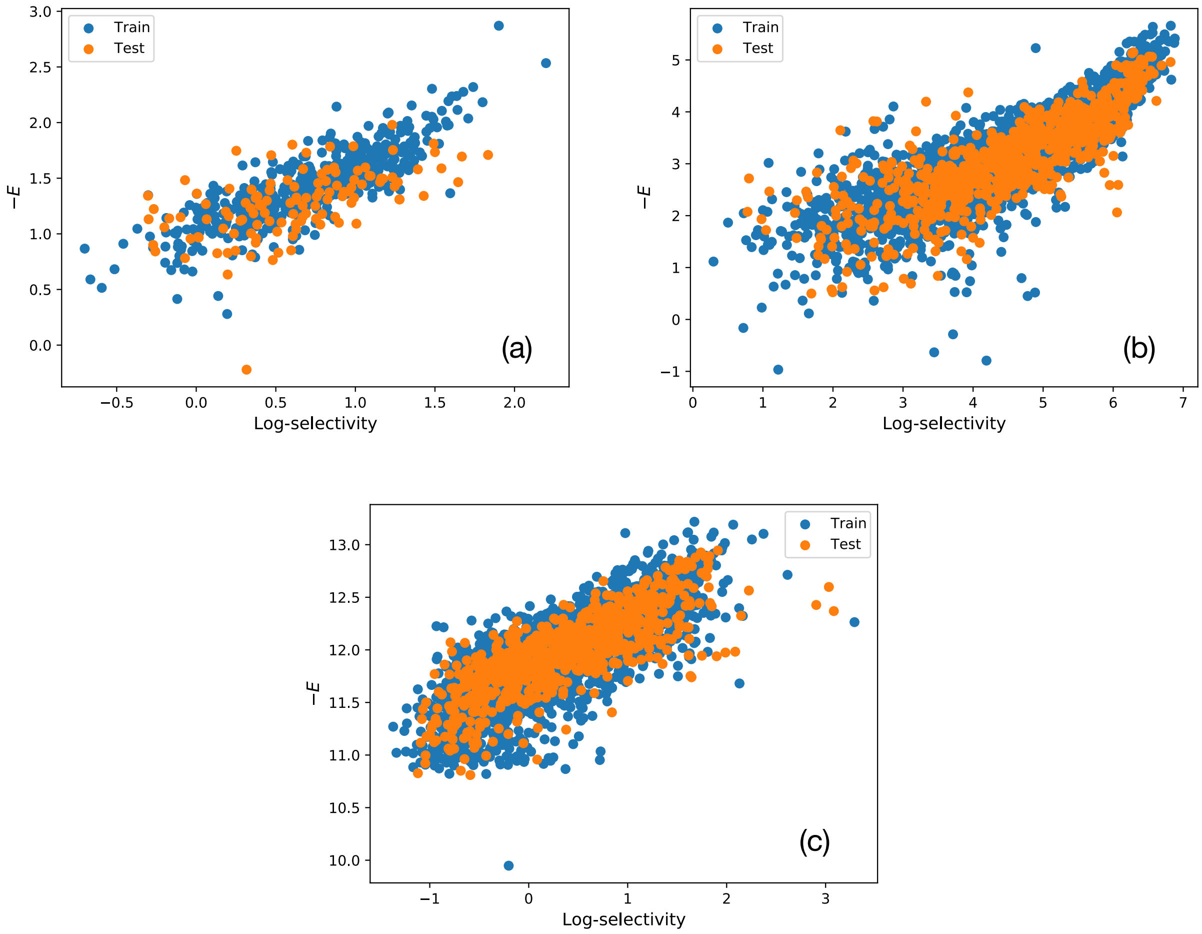


FIG. 2. Scatter plots between empirical log-selectivities and minus selective energies $E$. All the panels are realized retaining 1/25 of the total data points. Panel (a): scatter plot related to the dataset of [10], with Pearson correlation $\rho = 0.84$. Panel (b): scatter plot for the experiment [8], for which $\rho = 0.81$. Panel (c): scatter plot obtained from [9] data. The Pearson correlation is $\rho = 0.77$.

tion and Evolutionary Landscape by In Vitro Evolution, Molecular Biology and Evolution **37**, 1179 (2019), https://academic.oup.com/mbe/article-pdf/37/4/1179/32960043/msz256.pdf.

[4] E. Firnberg, J. W. Labonte, J. J. Gray, and M. Ostermeier, A comprehensive, high-resolution map of a gene's fitness landscape, Molecular biology and evolution **31**, 1581 (2014).

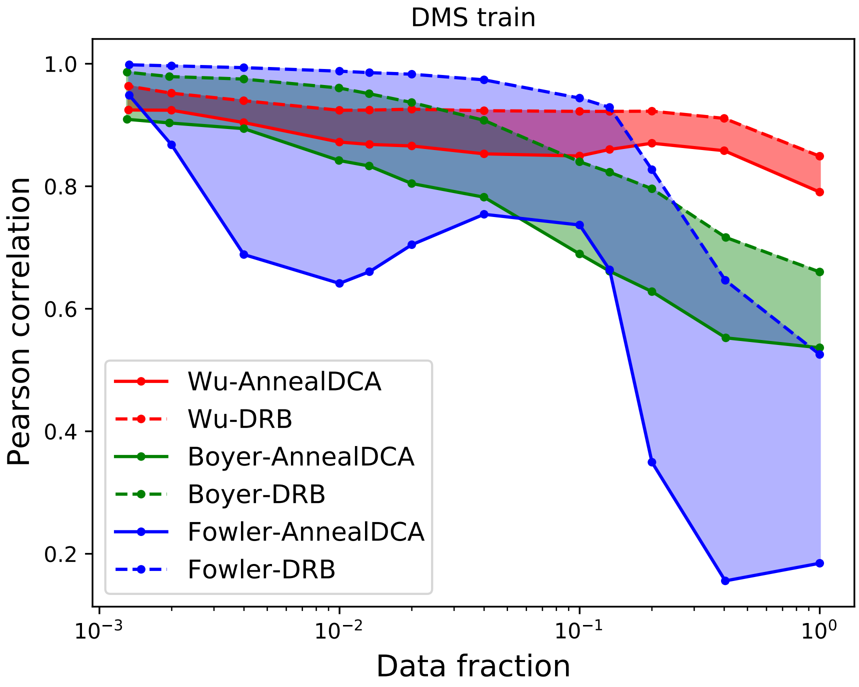


FIG. 3. Comparison between AnnealDCA and DRB [13] in terms of Pearson correlation between the inferred selective energies $E$ and the empirical log-selectivities, computed over the sequence in the train dataset. The correlation is reported as a function of the fraction of retained sequences, so to progressively exclude the ones characterized by a high uncertainty on log-selectivity.

[5] A. Gérard, A. Woolfe, G. Mottet, M. Reichen, C. Castrillon, V. Menrath, S. Ellouze, A. Poitou, R. Doineau, L. Briseno-Roa, *et al.*, High-throughput single-cell activity-based screening and sequencing of antibodies using droplet microfluidics, Nature biotechnology **38**, 715 (2020).

[6] T. A. Khan, S. Friedensohn, A. R. Gorter de Vries, J. Straszewski, H.-J. Ruscheweyh, and S. T. Reddy, Accurate and predictive antibody repertoire profiling by molecular amplification fingerprinting, Science advances **2**, e1501371 (2016).

[7] J. Dunbar and C. M. Deane, Anarci: antigen receptor numbering and receptor classification, Bioinformatics **32**, 298 (2016).

[8] N. C. Wu, L. Dai, C. A. Olson, J. O. Lloyd-Smith, and R. Sun, Adaptation in protein fitness landscapes is facilitated by indirect paths, Elife **5**, e16965 (2016).

[9] D. M. Fowler, C. L. Araya, S. J. Fleishman, E. H. Kellogg, J. J. Stephany, D. Baker, and S. Fields, High-resolution mapping of protein sequence-function relationships, Nature methods

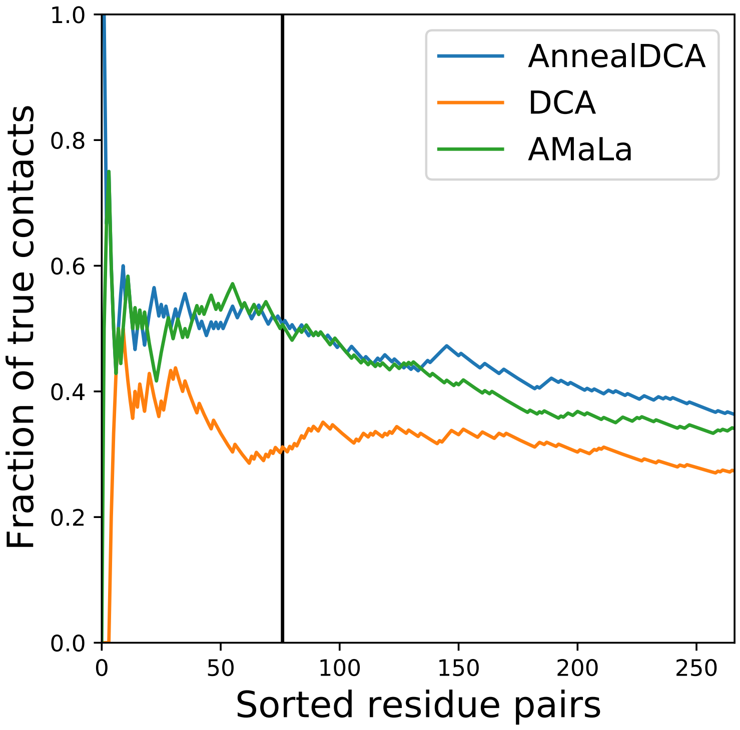


FIG. 4. Sensitivity plots for AAC6 contact prediction obtained with AnnealDCA, AMaLa and PlmDCA. On the horizontal axis the possible residue pairs are reported in a decreasing order of interaction score Eq. (5). On the vertical axis the fraction of correctly predicted contacts is reported. A solid vertical line is shown in correspondence of half the protein length.

**7**, 741 (2010).

[10] S. Boyer, D. Biswas, A. Kumar Soshee, N. Scaramozzino, C. Nizak, and O. Rivoire, Hierarchy and extremes in selections from pools of randomized proteins, Proceedings of the National Academy of Sciences **113**, 3482 (2016), https://www.pnas.org/content/113/13/3482.full.pdf.

[11] M. A. Stiffler, F. J. Poelwijk, K. P. Brock, R. R. Stein, A. Riesselman, J. Teyra, S. S. Sidhu, D. S. Marks, N. P. Gauthier, and C. Sander, Protein structure from experimental evolution, Cell Systems **10**, 15 (2020).

[12] D. M. Fowler and S. Fields, Deep mutational scanning: a new style of protein science, Nature methods **11**, 801 (2014).

[13] J. Fernandez-de Cossio-Diaz, G. Uguzzoni, and A. Pagnani, Unsupervised Inference of Protein Fitness Landscape from Deep Mutational Scan, Molecular Biology and Evolution

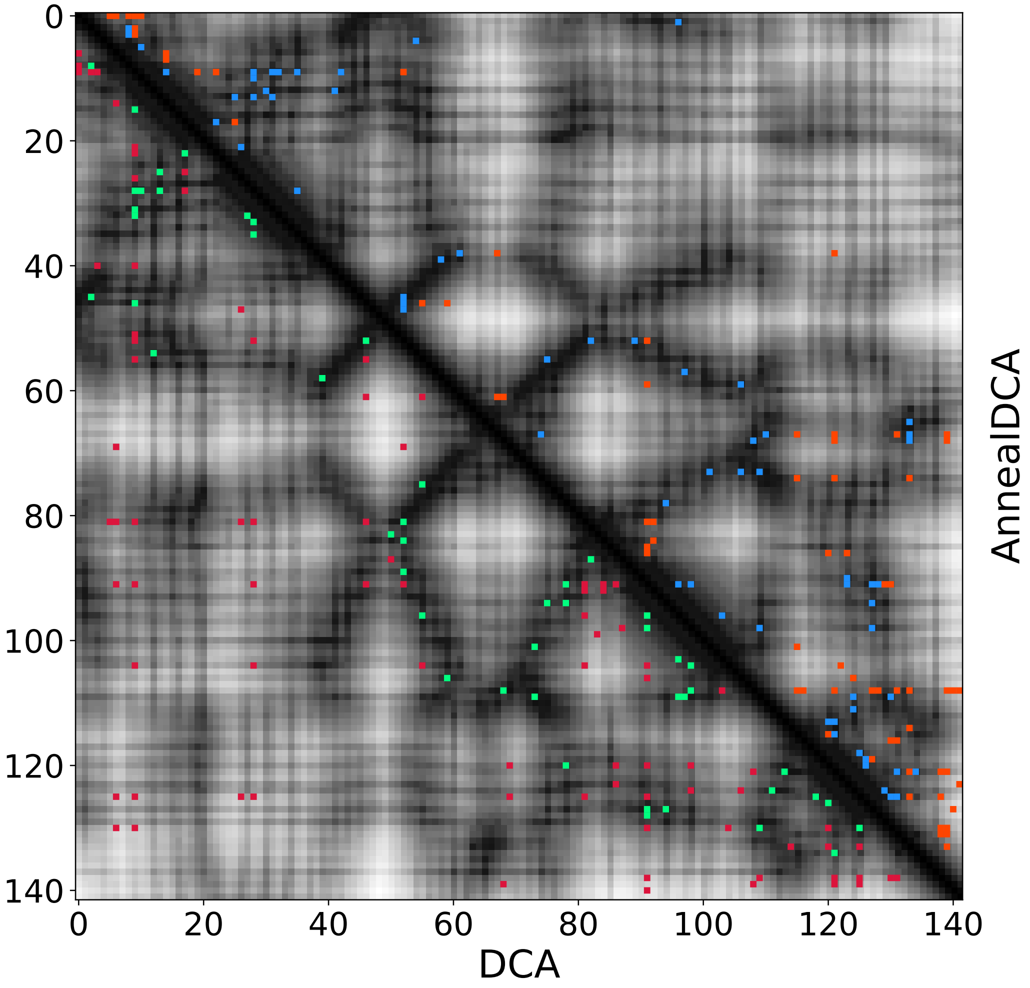


FIG. 5. Contact map prediction of AAC6 protein obtained with AnnealDCA (upper-right) and PlmDCA (lower-left). Correctly predicted contacts are reported in blue for AnnealDCA and green for PlmDCA, whereas wrong predictions are reported in orange for AnnealDCA and red for PlmDCA.

10.1093/molbev/msaa204 (2020), msaa204, https://academic.oup.com/mbe/advance-article-pdf/doi/10.1093/molbev/msaa204/33862547/msaa204.pdf.

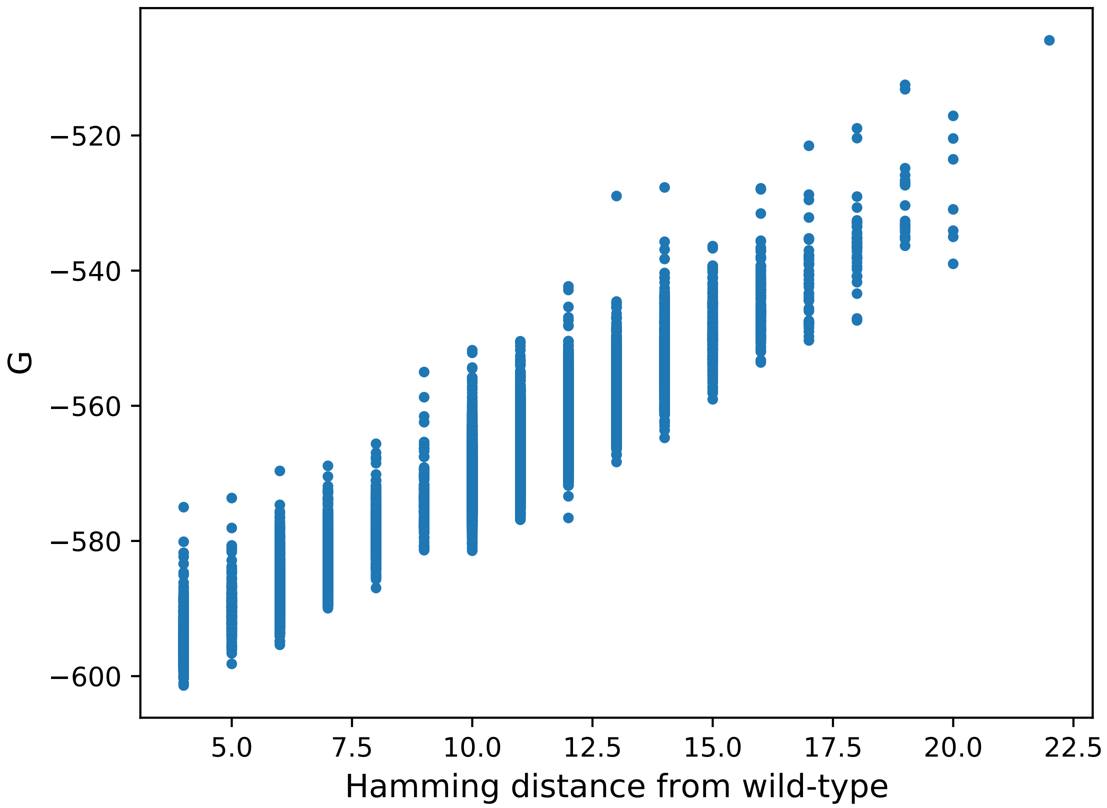


FIG. 6. Scatter plot between model energies $G$ computed over a set of sequences from AAC6 experiment and the Hamming distances of those sequences from the original wild-type.
